# Progressive excitability changes in the medial entorhinal cortex in the 3xTg mouse model of Alzheimer’s disease pathology

**DOI:** 10.1101/2023.05.30.542838

**Authors:** Lingxuan Chen, Zoé Christenson Wick, Lauren M. Vetere, Nick Vaughan, Albert Jurkowski, Angelina Galas, Keziah S. Diego, Paul Philipsberg, Denise J. Cai, Tristan Shuman

**Affiliations:** Icahn School of Medicine at Mount Sinai, New York NY; University of California Irvine, Irvine CA; CUNY Hunter College, New York NY; New York University, New York NY

## Abstract

Alzheimer’s disease (AD) is a chronic neurodegenerative disorder that is characterized by memory loss and progressive cognitive impairments. In mouse models of AD pathology, studies have found neuronal and synaptic deficits in the hippocampus, but less is known about what happens in the medial entorhinal cortex (MEC), which is the primary spatial input to the hippocampus and an early site of AD pathology. Here, we measured the neuronal intrinsic excitability and synaptic activity in MEC layer II (MECII) stellate cells, MECII pyramidal cells, and MEC layer III (MECIII) excitatory neurons at early (3 months) and late (10 months) time points in the 3xTg mouse model of AD pathology. At 3 months of age, prior to the onset of memory impairments, we found early hyperexcitability in MECII stellate and pyramidal cells’ intrinsic properties, but this was balanced by a relative reduction in synaptic excitation (E) compared to inhibition (I), suggesting intact homeostatic mechanisms regulating activity in MECII. Conversely, MECIII neurons had reduced intrinsic excitability at this early time point with no change in the synaptic E/I ratio. By 10 months of age, after the onset of memory deficits, neuronal excitability of MECII pyramidal cells and MECIII excitatory neurons was largely normalized in 3xTg mice. However, MECII stellate cells remained hyperexcitable and this was further exacerbated by an increased synaptic E/I ratio. This observed combination of increased intrinsically and synaptically generated excitability suggests a breakdown in homeostatic mechanisms specifically in MECII stellate cells at this post-symptomatic time point. Together, these data suggest that the breakdown in homeostatic excitability mechanisms in MECII stellate cells may contribute to the emergence of memory deficits in AD.

## INTRODUCTION

Alzheimer’s disease (AD) is an age-related neurodegenerative disorder that is characterized by memory loss and progressive cognitive impairments. While AD is defined by the presence of amyloid and tau pathology, the causal influence of these pathologies on the development and maintenance of cognitive impairments is poorly understood. Recent evidence suggests that reducing amyloid accumulations early in the disease may slow cognitive decline (van Dyck et al., 2023), but removing this pathology is not sufficient to fully rescue cognitive impairment (Musiek and Bennett, 2021; Karran and De Strooper, 2022; van Dyck et al., 2023). This suggests that understanding the early circuit changes that drive the onset of memory impairments will be key to early AD detection and intervention. In particular, region-specific changes in excitability can precede the buildup of aggregated amyloid (Buckner et al., 2005), suggesting that early changes in cellular and circuit excitability may contribute to cognitive deficits. It is critical, then, to understand the specific cellular and circuit properties that are altered early in disease progression, even before the onset of cognitive deficits.

The entorhinal cortex is the primary spatial input to the hippocampus and has long been implicated as an early site of AD pathology and cell loss in humans (Hyman et al., 1986; Braak and Braak, 1991; Gómez-Isla et al., 1996; Masdeu et al., 2005; Stranahan and Mattson, 2010). This suggests that early damage to the entorhinal-hippocampal system may drive early cognitive deficits in AD. Indeed, the first symptoms observed in patients with AD are often spatial and episodic memory deficits (Burgess et al., 2002). These findings are also recapitulated in mouse models of AD pathology, where studies have found impairments in hippocampus-dependent spatial memory (Billings et al., 2005; Saito et al., 2014) and spatial coding (Cacucci et al., 2008; Mably et al., 2017; Jun et al., 2020). Accordingly, hippocampal deficits in neuronal excitability (Ohno et al., 2004; Brown et al., 2011; Kaczorowski et al., 2011; Davis et al., 2014; Kerrigan et al., 2014; Frazzini et al., 2016) and synaptic transmission (Oddo et al., 2003; Palop and Mucke, 2010; Booth et al., 2016b; Forner et al., 2017; Chen et al., 2018) have been observed across various mouse models. In addition, recent evidence suggests that spatial coding in the medial entorhinal cortex (MEC) is disrupted in mouse models of AD pathology, with deficits in grid cell coding observed (Fu et al., 2017; Ying et al., 2022; Igarashi, 2023) even before the onset of spatial memory impairment (Jun et al., 2020). Similarly, humans at genetic risk for AD also have reduced grid-like representations in the temporal lobe that were present before to the emergence of spatial memory deficits (Kunz et al., 2015). It is therefore likely that deficits in MEC grid representations are contributing to the decline in spatial coding and spatial memory seen across AD patients and rodent models of AD pathology. However, the specific cellular and circuit processes that break down in the MEC to drive altered spatial coding in AD remain unknown.

In MEC layer 2 (MECII), there are 2 major types of excitatory cells, stellate and pyramidal cells, that have distinct electrophysiological properties (Alonso and Klink, 1993; Canto and Witter, 2012; Fuchs et al., 2016). Conversely, in MEC layer 3 (MECIII), the excitatory cells are generally homogenous, with similar intrinsic properties to MECII pyramidal cells (Dickson et al., 1997; Gloveli et al., 1997). Importantly, these three excitatory cell types in superficial MEC have distinct major anatomical projections. MECII stellate cells primarily project to the dentate gyrus, forming the origin of the tri-synaptic pathway, while MECIII excitatory cells provide the primary direct spatial input to CA1 through the temporoammonic pathway (Suh et al., 2011; Kitamura et al., 2014; Witter et al., 2014). Finally, MECII pyramidal neurons primarily provide local and commissural projections within MEC (Ohara et al., 2019), as well as some input directly to CA1 (Kitamura et al., 2014; Ohara et al., 2019). Within these distinct subpopulations, there is evidence that the intrinsic properties of MECII stellate cells may be specifically affected in rodent models of AD pathology (Stranahan and Mattson, 2010; Marcantoni et al., 2014; Booth et al., 2016a; Heggland et al., 2019). However, the field lacks a thorough examination of each of these populations across the progression of memory impairments, and thus, it is important to characterize the nature and timing of dysfunction in these distinct subtypes of MEC excitatory neurons.

Neuronal intrinsic excitability is an important cellular property that reflects a neuron’s tendency to generate action potentials upon synaptic integration, and is critical for learning and memory processes (Daoudal and Debanne, 2003; Zhang and Linden, 2003; Sehgal et al., 2013; Chen et al., 2020). While there are relatively few studies looking at AD-related changes in neuronal intrinsic excitability in MEC, there is some evidence that MECII stellate cells may be hyperexcitable in mouse models of AD pathology (Stranahan and Mattson, 2010; Marcantoni et al., 2014; Booth et al., 2016a; Heggland et al., 2019). Notably, driving increased excitability in MECII stellate cells with chemogenetics can lead to poor hippocampal CA1 spatial coding, place cell remapping, and memory deficits in wild-type mice (Kanter et al., 2017). This suggests that MECII stellate cell hyperexcitability is sufficient to drive cognitive dysfunction and may contribute to memory deficits in AD. In addition to a neuron’s intrinsic properties, its activity is also tuned by the amount of excitatory and inhibitory synaptic inputs onto the cell. In mouse models of AD pathology, there is evidence of interneuron dysfunction that has been shown to lead to aberrant network excitation and memory impairment (Verret et al., 2012; Palop and Mucke, 2016; Schmid et al., 2016). Therefore, besides neuronal intrinsic excitability, it is crucial to probe synaptic excitation/inhibition (E/I) balance in the same neurons to get a more thorough assessment of the network state in the healthy and diseased brain.

Here, we followed intrinsically and synaptically generated excitability in the MEC across disease progression using the 3xTg mouse model of AD pathology which has both amyloid and tau mutations as well as progressive spatial memory impairments (Oddo et al., 2003; Billings et al., 2005; Javonillo et al., 2022). We performed *in vitro* whole-cell patch clamp recordings in MECII stellate cells, MECII pyramidal cells, and MECIII excitatory cells in WT and 3xTg mice at early (3 months; before the onset of spatial memory deficits, referred to as pre-symptomatic) and late (10 months; after spatial memory deficits have emerged, referred to as post-symptomatic) time points and measured both neuronal intrinsic excitability and spontaneous synaptic activity. At the pre-symptomatic time point, we found changes in excitability in all cell types, but hyperexcitability in MECII stellate and pyramidal cells was balanced by corresponding shifts in the E/I ratio of synaptic inputs, suggesting strong homeostatic control of overall activity. By the late, post-symptomatic time point, MECII stellate cells showed increases in both intrinsic excitability and synaptic E/I ratio, while MECII and MECIII pyramidal cells were mostly normalized. These results suggest that a breakdown in homeostasis drives increased excitability in MECII stellate cells that may contribute to the progression of memory impairments in 3xTg mice.

## MATERIALS AND METHODS

### Animals

All experiments were approved in advance by the Institutional Care and Use Committee of the Icahn School of Medicine at Mount Sinai. Both male and female transgenic 3xTg (MMRRC 034830-JAX) (Oddo et al., 2003) and wild-type (WT) (B6129SF2/J, JAX 101045) mice (Gstir et al., 2014; Stevens and Brown, 2015) were used. Breeders were purchased from the Mutant Mouse Resource and Research Center (MMRRC) and the colony was maintained in approved breeding facilities at Mount Sinai. Mice were given ad libitum food and water on a 12-hour light-dark cycle (lights on at 0700 hours).

### Novel Object Location

Prior to all experiments, mice were habituated to transportation and handled in the behavior room for 5 days. Mice were then habituated to an empty arena (1ft x 1ft) with spatial cues on the walls for 5 minutes over 2 consecutive days. The next day, mice were returned to the arena and explored 2 identical objects (plastic toys ∼2 × 2 inch) taped near two corners of the arena (training session; **Fig. 1A**). Mice were then returned to their home cage for 1 or 4 hours before testing. During testing, one of the objects was moved to a new location within the arena and the subject was allowed to freely explore both objects (test session; **Fig. 1A**). To quantify exploration, we recorded each session with an overhead webcam running Miniscope recording software (Cai et al., 2016; Shuman et al., 2020) and scored interaction bouts using Chronotate, a customized Python application that timestamps user inputs during video playback (available at https://github.com/ShumanLab/Chronotate). Interaction was defined as active investigation of the objects, and only the first 15s of interaction during the test session was used. Mice that had a strong preference for one object (DI>0.25 or DI<-0.25) during the training sessions or exclusively interacted with one object during the test sessions were excluded. Discrimination index (DI) was calculated as:

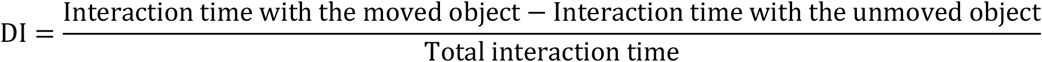

Note that we used two slightly different protocols for this experiment. For the “4hr” group, mice spent 10 minutes in the arenas to explore the objects (training session), then 4 hours later, spent 10 minutes in the same arena for the test session. For the “1hr” group, the delay between training and test was 1 hour and during both training and test, an experimenter was counting mice’s interaction with both objects in real time with a stopwatch. Once mice reached either 30 s of total interaction or 20 min in the box they were taken out. We observed no difference in the DI of test sessions between these two protocols (**Table 1**) and the trends from all groups were nearly identical. Thus, we have collapsed across these two delay periods for behavioral analysis.

**Figure 1.**
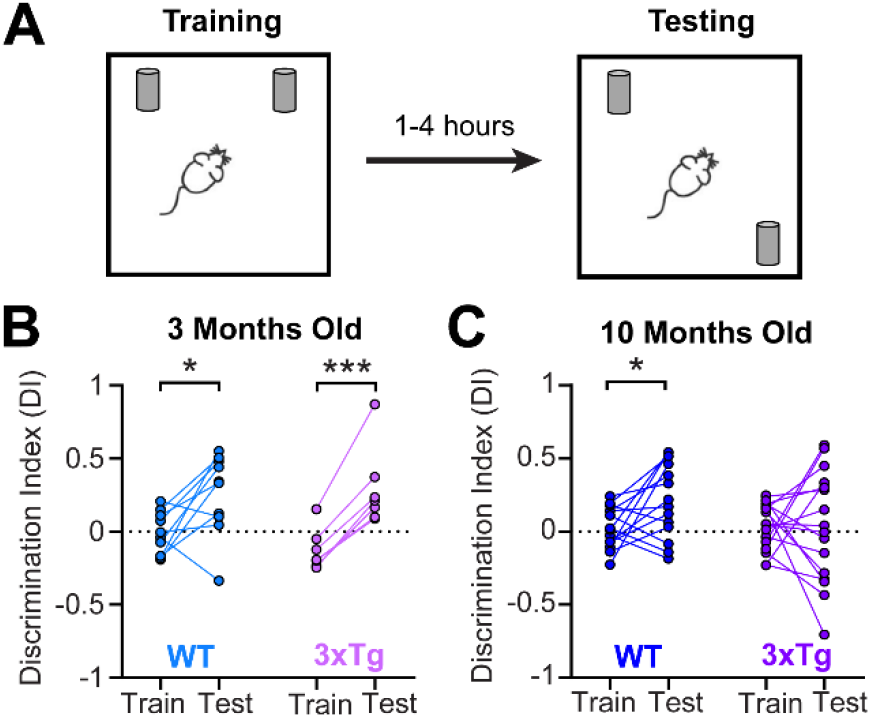
3xTg mice had intact spatial memory in the novel object location task at 3 months of age but developed a deficit by 10 months of age. (**A**) Schematic of the novel object location task. (**B**) At 3 months of age, both WT and 3xTg mice showed a preference for the newly located object during the test session, indicated by a significantly increased Discrimination Index (DI) in the test session compared to the training session (paired t-test from Training to Testing, WT: *p*=0.023, 3xTg: *p*<0.001). (**C**) At 10 months of age, while WT mice still showed a preference for the moved object in the test session (paired t-test from Training to Testing, WT: *p*=0.037), 3xTg mice did not (paired t-test from Training to Testing, 3xTg: *p*>0.05), indicating impaired memory at this late time point.

**Table 1.** Test session discrimination index (DI) in the novel object location task with the 1hr and 4hr protocols in WT and 3xTg mice at 3m and 10m.

| Group | 1hr | 4hr | p value | Test |
| --- | --- | --- | --- | --- |
| 3m WT | Values: 0.34, 0.48, -0.34, 0.04, 0.55, 0.12<br>Mean±SEM: 0.20 ± 0.13 | Values: 0.44, 0.50, 0.10, 0.33<br>Mean±SEM: 0.34 ± 0.09 | 0.4522 | Unpaired t-test |
| 3m 3xTg | Values: 0.87, 0.16, 0.37, 0.21, 0.09<br>Mean±SEM: 0.34 ± 0.14 | Values: 0.23, 0.10<br>Mean±SEM: 0.17 ± 0.07 | 0.4917 | Unpaired t-test |
| 10m WT | Values: 0.33, -0.19, 0.54, 0.17, -0.14<br>Mean±SEM: 0.25 ± 0.12 | Values: 0.06, 0.52, 0.22, -0.08, 0.38, 0.03, 0.12<br>Mean±SEM: 0.18 ± 0.08 | 0.6412 | Unpaired t-test |
| 10m 3xTg | Values: 0.04, 0.28, -0.15, 0.30, -0.44, -0.33<br>Mean±SEM: 0.00 ± 0.10 | Values: 0.15, -0.28, -0.71, 0.59, 0.00, 0.57, 0.45, -0.34<br>Mean±SEM: 0.05 ± 0.17 | 0.7935 | Unpaired t-test |

### In vitro whole-cell patch clamp recordings

Mice were anesthetized by isoflurane through inhalation, followed by rapid decapitation. Horizontal acute brain slices were prepared on a vibratome (Leica VT1200S, Leica Biosystems, IL) at 300 μm thickness in ice-cold cutting solution (in mM: 135 NMDG, 10 D-glucose, 4 MgCl_2_, 0.5 CaCl_2_, 1 KCl, 1.2 KH_2_PO_4_, 20 HEPES, and sucrose to adjust the osmolality to 305–310 mmol/kg; pH 7.35; bubbled with 95% O_2_/5% CO_2_). Slices were kept in a submerged chamber filled with sucrose-based artificial cerebrospinal fluid (sACSF; in mM: 55 sucrose, 85 NaCl, 25 D-glucose, 2.5 KCl, 1.25 NaH_2_PO_4_, 0.5 CaCl_2_, 4 MgCl_2_, 26 NaHCO_3_; 300–305 mmol/kg; bubbled with 95%O_2_/5% CO_2_). The recovery chamber was kept in 37°C water bath for 30 min before being held at room temperature.

For whole-cell patch clamp recordings, acute brain slices were placed in a submerged chamber filled with circulating room temperature ACSF (in mM: 126 NaCl, 10 D-glucose, 2 MgCl_2_, 2 CaCl_2_, 2.5 KCl, 1.25 NaH_2_PO_4_, 1.5 sodium pyruvate, 1 L-glutamine; 295–300 mmol/kg; bubbled with 95%O_2_/5% CO_2_). A K-met based intracellular solution (in mM: 127.5 K-methanesulfonate, 10 HEPES, 5 KCl, 5 Na-phosphocreatine, 2 MgCl_2_, 2 Mg-ATP, 0.6 EGTA, and 0.3 Na-GTP; pH 7.25; 285-300 mOsm) was used for the current- and voltage-clamp recordings to probe neuronal intrinsic excitability and spontaneous postsynaptic currents, respectively.

Cells were visualized on a Nikon Eclipse FN1 microscope (Nikon Instrument Inc., NY) paired with a SOLA light engine (Lumencor, OR) to identify cells in MEC layers II or III. A Multiclamp 700B amplifier and an Axon Digidata 1550B digitizer (Molecular Devices Inc., CA) were used for data acquisition. Whole-cell recordings were achieved under voltage-clamp at -60 mV. Cells were next switched to current clamp with I=0. Resting membrane potential was measured when cells stabilized. Current-spike curves were determined by injecting a series of current steps (0-500 pA at 50 pA increments, 500 ms) under I=0. A series of other intrinsic excitability measurements were done while cells were held at -70 mV: 1) Rheobase, calculated as the current needed to generate the first action potential, by injecting a series of 500 ms current steps at 10 pA increments; 2) Input resistance, calculated as Rin = V / I, at I=-50 pA; 3) Spike amplitude: amplitude of the first stimulated spike; 4) Spike half-width: width of the first stimulated spike at its half maximal value; 5) Medium afterhyperpolarization (mAHP), calculated as the lowest point of the post-burst hyperpolarization relative to baseline, averaged over five 500 ms, 500 pA current steps; 6) Slow afterhyperpolarization (sAHP), calculated as the remaining hyperpolarization 1.5 s after the end of stimulation, averaged over five 500 ms, 500 pA current steps; 7) Sag ratio, the amount of sag potential during a hyperpolarizing current step (I=-100 pA), calculated as *(lowest point of hyperpolarization – steady state voltage at the end of current step) / (lowest point of hyperpolarization – pre-step baseline)*. After these measurements, cells were switched back to voltage clamp at V=-60 mV to record 90 s of spontaneous EPSCs, then slowly depolarized to V=0 to record 90 s of spontaneous IPSCs. Clampfit 10.7 (Molecular Devices Inc., CA) was used for processing electrophysiology data. Action potential measurements were done using the “Threshold Search” mode under “Event Detection”. Recordings acquired under voltage clamp were first filtered with a Gaussian low-pass filter. Then, EPSCs and IPSCs were detected using the “Template Search” mode, using cell-based templates. Detected events also went through a visual inspection to exclude any outliers or noise.

### Experimental Design and Statistical Analyses

Both male and female mice were used and the number of mice of each sex used for behavior and patch clamp recordings and the total number of cells recorded are listed in **Table 2**. For each experiment, the number of cells (*n*) and number of animals (*N*) are given in the corresponding figure legend. To determine the progression of memory deficits in the 3xTg model for the novel object location task (**Fig. 1**), mice at 2.3-4.1 months and 9.2-11.5 months of age were used for group “3m” and “10m”, respectively. To examine intrinsic and synaptic excitability at the early and late time points, mice at 3.3-4.0 months and 9.0-11.6 months of age were used for group “3m” and “10m”, respectively, for whole-cell patch clamp recordings.

**Table 2.** Number of animals (N) of each sex and number of recorded cells (n).

| Novel object location (1hr delay) | 3 m WT | 3 m 3xTg | 10 m WT | 10 m 3xTg |
| --- | --- | --- | --- | --- |
| N of males | 3 | 3 | 4 | 6 |
| N of females | 3 | 2 | 3 | 2 |
| Novel object location (4hr delay) | 3 m WT | 3 m 3xTg | 10 m WT | 10 m 3xTg |
| N of males | 2 | 0 | 3 | 3 |
| N of females | 2 | 2 | 4 | 5 |

**Table 2.** Number of animals (N) of each sex and number of recorded cells (n).
| MECII recordings | 3 m WT | 3 m 3xTg | 10 m WT | 10 m 3xTg |
| --- | --- | --- | --- | --- |
| N of males | 1 | 3 | 2 | 1 |
| n of males | 1 | 15 | 2 | 2 |
| n (stellate) of males | 0 | 8 | 0 | 2 |
| n (pyramidal) of males | 1 | 7 | 2 | 0 |
| N of females | 9 | 5 | 7 | 6 |
| n of females | 29 | 19 | 30 | 24 |
| n (stellate) of females | 7 | 13 | 11 | 14 |
| n (pyramidal) of females | 22 | 6 | 19 | 10 |
| MECIII recordings | 3 m WT | 3 m 3xTg | 10 m WT | 10 m 3xTg |
| N of males | 0 | 0 | 3 | 0 |
| n of males | 0 | 0 | 9 | 0 |
| N of females | 4 | 3 | 1 | 4 |
| n of females | 12 | 13 | 1 | 10 |

#### Clustering MECII neurons into stellate and pyramidal cells

During whole-cell patch clamp recordings in layer II of MEC, experimenters provided a preliminary identification of whether a cell was stellate or pyramidal by its morphology under the microscope and the action potential firing pattern. To provide an unbiased approach, we clustered MECII neurons into two subtypes based on electrophysiological parameters. Previous literature has indicated significantly larger hyperpolarizing sag currents (Alonso and Klink, 1993; Canto and Witter, 2012; Fuchs et al., 2016; Fernandez et al., 2022) and AHPs (Alonso and Klink, 1993) in stellate cells compared to pyramidal cells in MECII. Therefore, we used k-means clustering to divide cells into two sets based on the “sag ratio” and mAHP (see “*In vitro whole-cell patch clamp recordings*”), which gave clear separation between clusters (**Fig. 2**). To accommodate potential age-related changes, we performed the k-means clustering on 3-month (**Fig. 2A**) and 10-month (**Fig. 2B**) data separately. Note that WT and 3xTg cells at each time point were clustered together, but for the purpose of visualization they were separately plotted. We also visually inspected the automated clustering to ensure proper performance of the algorithm. Based on visual inspection of the clusters plus the preliminary identification of cell type during patching, we manually changed the cluster of 1 cell in the 3-month 3xTg group and 1 cell in the 10-month WT group and excluded 2 cells in the 3-month WT group that lay on the border line between clusters. These changes were made while blind to all other excitability measurements. In addition, there were 6 cells in the 3-month 3xTg group and 5 cells in the 10-month 3xTg group that did not have mAHP measurements. These 11 cells are not shown in **Fig. 2**. They were only manually clustered as stellate cells if: 1) sag ratio was greater than 0.3, and 2) during recording they were visually determined to be stellate cells. Otherwise, they were clustered as pyramidal cells.

**Figure 2.**
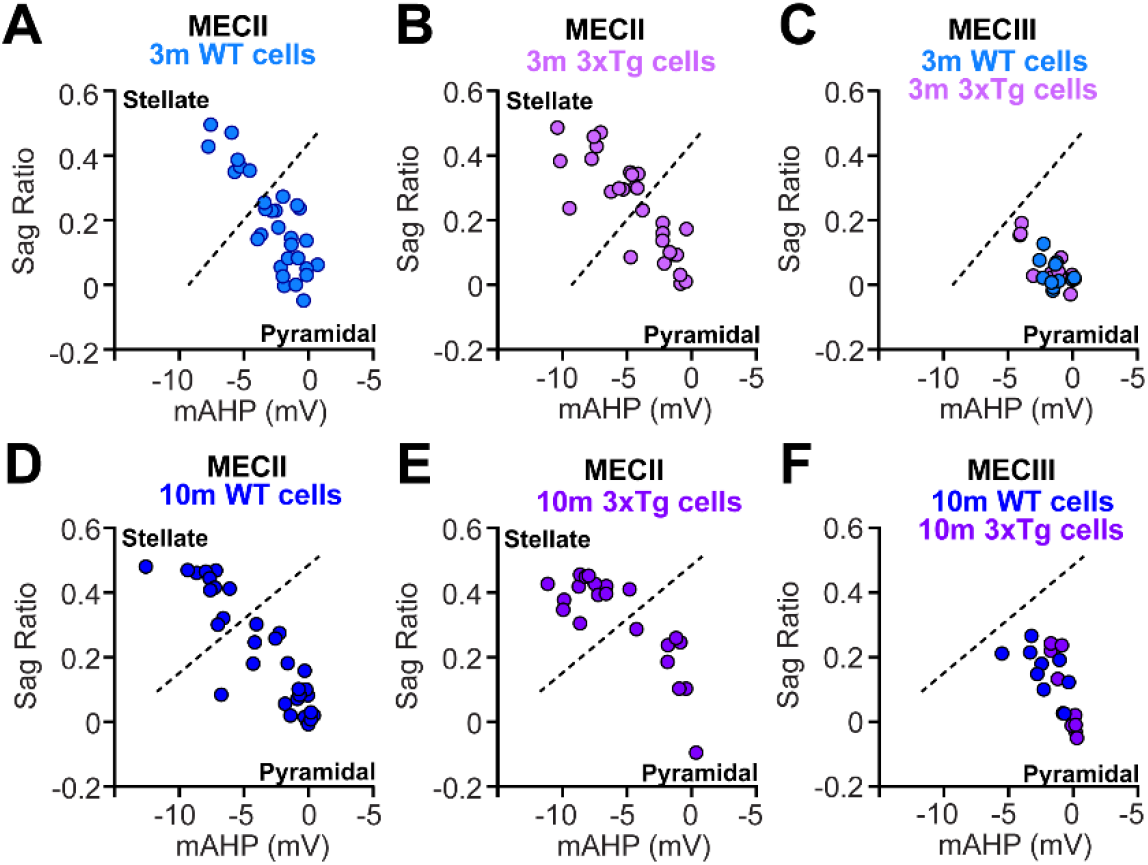
MECII neurons were clustered into stellate- and pyramidal-like cells based on their electrophysiological properties, while MECIII neurons had properties similar to pyramidal cells. (**A, B, D, E**) Using k-means clustering, MECII neurons recorded at 3 months (**A**, n=30 cells from 10 WT mice, **B**, n=28 cells from 7 3xTg mice) and 10 months (**D**, n=32 cells from 9 WT mice, **E**, n=21 from 5 3xTg mice) of age were separated into two clusters based on the sag ratio and mAHP of each neuron. Neurons with large sag ratios and mAHPs were clustered as stellate-like cells (above the dashed lines), and those with small sag ratios and mAHPs were clustered as pyramidal cells (below the dashed lines). Note that WT and 3xTg cells were clustered together, but for the purpose of visualization they were separately plotted. (**C, F**) 3-month (**C**, n=12 cells from 4 WT mice, n=13 from 3 3xTg mice) and 10-month (**F**, n=9 from 4 WT mice, n=10 from 4 3xTg mice) MECIII cells recorded from WT and 3xTg mice all fell into the pyramidal cell cluster when using the same criteria as layer II data (A, B, D, E).

#### Data analysis and statistics of patch-clamp recordings

Current-evoked action potentials under current-clamp and postsynaptic currents under voltage-clamp were detected using Clampfit 10.7 (Molecular Devices Inc., CA) plus a visual inspection (see “*In vitro whole-cell patch clamp recordings*”). Statistical analyses were performed with GraphPad Prism 9 (GraphPad Software, CA). For all analyses, outliers in each data group were first identified with the ROUT method, Q=1% and excluded from the dataset. Then, normality was determined using Shapiro-Wilk test. An unpaired t-test was used for normally distributed data and a Mann–Whitney test was used for data that was not normally distributed. For all graphs, data are represented as mean ± SEM (standard error of mean). For comparisons between the current-spike curves, cells that did not receive all the current steps from 0-500 pA (for MECII neurons) or from 0-300 pA (for MECIII neurons) were excluded, and two-way repeated measures ANOVA (RM ANOVA) was used to test for statistical differences between genotypes.

## RESULTS

### Impaired spatial memory in 10-month, but not 3-month-old, 3xTg mice

To examine the progression of spatial memory deficits in the 3xTg mouse model of AD pathology, we used a novel object location task (**Fig. 1A**) that is dependent on both hippocampus (Murai et al., 2007; Wimmer et al., 2012) and MEC (Tennant et al., 2018). During initial training, WT and 3xTg mice investigated two identical objects, showing no preference for one over another, indicated by a DI that was not significantly different from 0, at either 3 months (**Fig. 1B**, Training, one sample t-test, *p*>0.05 for both WT and 3xTg mice) or 10 months of age (**Fig. 1C**, Training, one sample t-test, *p*>0.05 for both WT and 3xTg mice). During testing, the animals were returned to the arena 1 or 4 hours later with one object moved to a new location (see **Materials and Methods** and **Tables 1 & 2** for details). In 3-month-old mice, both WT and 3xTg mice showed a preference for the newly located object (**Fig. 1B**, Test, one sample t-test, WT: *p*=0.016, N=10, 3xTg: *p*=0.030, N=7; paired t-test from Training to Testing, WT: *p*=0.023, 3xTg: *p*<0.001), indicating an intact spatial memory of the unmoved object. In contrast, in 10-month-old mice, only the WT mice showed significant learning during the test (**Fig. 1C**, Test, one sample t-test, WT: *p*=0.009, N=14; paired t-test from Training to Testing, WT: *p*=0.037). 10-month-old 3xTg mice did not show a preference for the moved object (**Fig. 1C**, Test, one sample t-test, 3xTg: *p*>0.05, N=16; paired t-test from Training to Testing, 3xTg: p>0.05), suggesting impaired spatial memory at this late time point.

Overall, these results show that spatial memory deficits are age-dependent and emerge between 3 and 10 months of age in the 3xTg mice, consistent with previous reports (Billings et al., 2005; Davis et al., 2013; Belfiore et al., 2019; Creighton et al., 2019). Therefore, we utilized 3- and 10-month-old mice for our pre- and post-symptomatic time points for all additional experiments.

### Classification of MECII cells into stellate and pyramidal cells in 3-month and 10-month-old WT and 3xTg mice based on electrophysiological properties

The cell types in medial entorhinal cortex layer II (MECII) and III (MECIII) have been well-characterized based on morphological, immunohistochemical, and electrophysiological evidence over the past few decades (Alonso and Klink, 1993; Klink and Alonso, 1997; Varga et al., 2010; Canto and Witter, 2012; Kitamura et al., 2014; Ray et al., 2014). In MECII, stellate cells have distinct intrinsic electrophysiological properties not present in MECII pyramidal cells, such as significant hyperpolarizing sag potentials, burst firing, and depolarizing afterpotentials (Alonso and Klink, 1993; Canto and Witter, 2012; Fuchs et al., 2016; Fernandez et al., 2022). This allowed us to identify MECII stellate versus pyramidal cells based on these intrinsic properties. We found that the sag ratio (at -100 pA current step) and medium afterhyperpolarizing potential (mAHP) amplitudes best isolated the clusters of MECII neurons in our recordings (**Fig. 2;** see **Materials and Methods**). In accordance with the literature, stellate cells were identified as the cluster with high sag potential and large mAHP (**Fig. 2A, B, D, E** above the dashed lines), and pyramidal cells were identified as the cluster with low sag ratio and small mAHP (below the dashed lines). To further validate our cell-type classification, we have plotted MECIII cells in the same space (**Fig. 2C, F**). Since layer III neurons have been shown to have properties similar to MECII pyramidal cells (Dickson et al., 1997; Gloveli et al., 1997), they should all fall into the pyramidal cluster. Indeed, all 3m and 10m MECIII cells fell below the dashed line (**Fig. 2C, F**). In addition to validating our MECII clustering, this also supported the identification of MECIII neurons as pyramidal cells. Together, this clustering strategy allowed us to identify MECII stellate cells, MECII pyramidal cells, and MECIII excitatory neurons based on their well-characterized intrinsic properties.

### Early changes in neuronal excitability are balanced by corresponding changes in synaptic inputs in MECII and MECIII cells of 3xTg mice at the pre-symptomatic time point

To test for changes in neuronal excitability at the early, pre-symptomatic time point, we performed whole-cell patch clamp recordings in the MEC of 3xTg and WT mice at 3 months of age. In MECII stellate cells of 3xTg mice (**Fig. 3**), we found evidence of enhanced intrinsic excitability at this early time point. While the resting membrane potential of 3xTg MECII stellate cells was comparable to that of WT cells (**Fig. 3A**, unpaired t-test, p>0.05), the rheobase, defined as the minimum current needed to elicit an action potential, of 3xTg cells was significantly reduced (**Fig. 3B**, unpaired t-test, *p*=0.009). Further, the number of evoked action potentials in response to current injections of increasing size was significantly increased in 3xTg cells (**Fig. 3C**, two-way RM ANOVA, Main Effect of Genotype: *p*=0.001). These results indicate that there is intrinsic hyperexcitability in MECII stellate cells in 3xTg mice at 3 months, before the onset of cognitive deficits. However, when we looked at spontaneous excitatory and inhibitory postsynaptic currents (EPSC and IPSC, respectively) in the same stellate cells, we found that the ratio of excitatory to inhibitory inputs (E/I ratios) was altered. Specifically, we found that both the ratio of E/I frequency and E/I amplitude were significantly skewed towards inhibition, suggesting relatively more robust inhibitory inputs to MECII stellate cells in 3xTg mice (**Fig. 3D-F**, E/I frequency, unpaired t-test, *p*=0.007; **Fig. 3G-I**, E/I amplitude, Mann-Whitney, *p*=0.031). In particular, the reduction in the ratio of E/I amplitude was primarily caused by increased IPSC amplitude in 3xTg mice (**Fig. 3H**, IPSC amplitude, Mann-Whitney, *p*=0.025). Taken together, these opposing changes to intrinsic and synaptic excitability, which are likely a combination of pathological and compensatory homeostatic changes, may result in little to no net change in neuronal activity in 3-month-old 3xTg mice.

**Figure 3.**
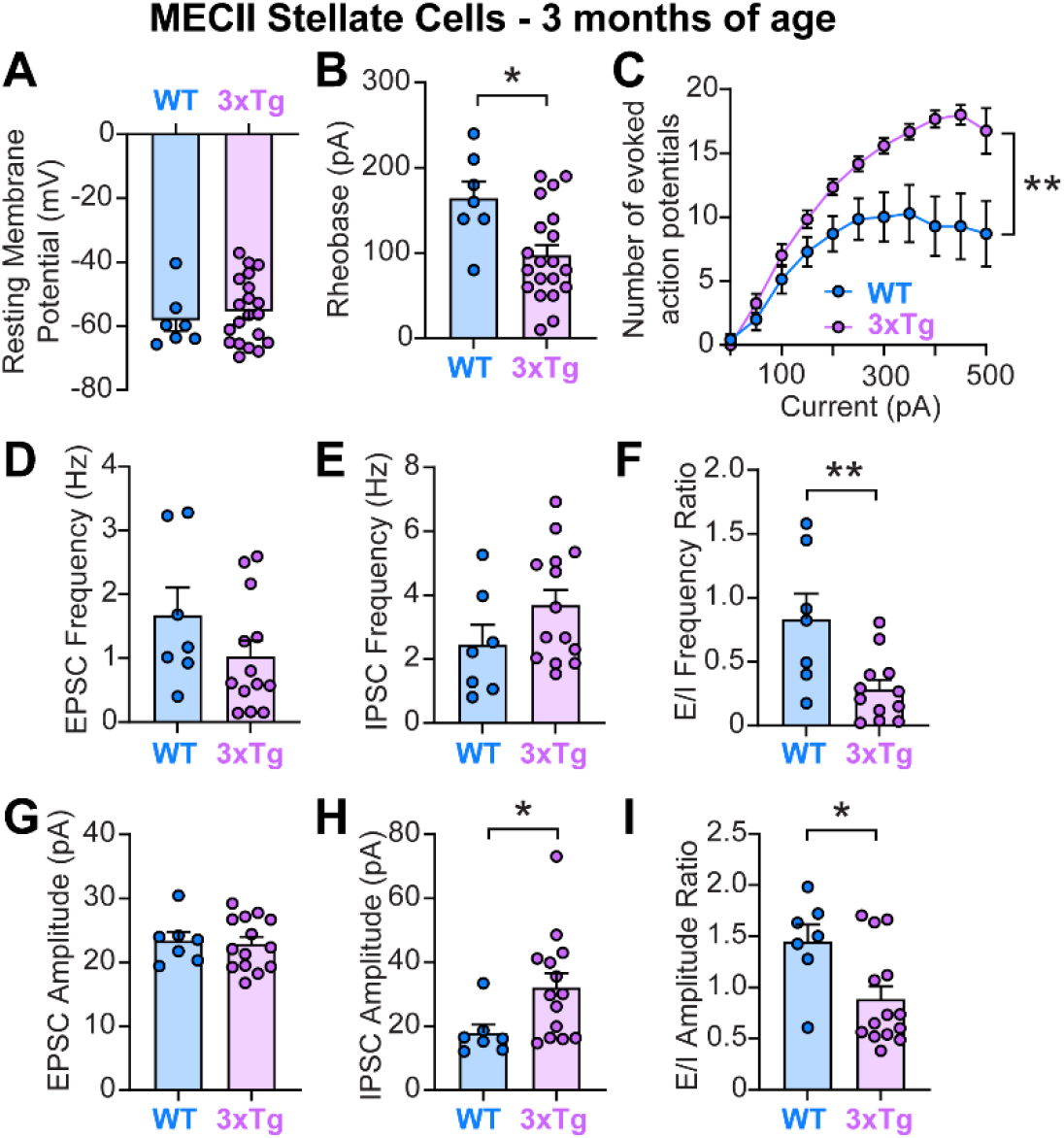
At 3 months of age, MECII stellate cells in 3xTg mice had increased neuronal intrinsic excitability, which was balanced by increased synaptic inhibition and decreased synaptic E/I ratio. (**A-C**) Neuronal intrinsic excitability of MECII stellate cells was increased in 3xTg mice compared to controls, indicated by a reduced rheobase (**B**, unpaired t-test, *p*=0.009) and an increased number of current-evoked action potentials (**C**, n=7 cells from WT mice; n=12 cells from 3xTg mice; two-way RM ANOVA, Main Effect of Genotype: *p*=0.001). (**D-I**) Meanwhile, there was increased synaptic inhibition onto these stellate cells, demonstrated by a larger IPSC amplitude (**H**, Mann-Whitney test, *p*=0.025) and an overall decrease in synaptic E/I ratio both in frequency (**F**, unpaired t-test, *p*=0.007) and amplitude (**I**, Mann-Whitney test, *p*=0.031). All other tests showed no differences between groups (*p*>0.05).

We next examined MECII pyramidal cells at the same early time point and found a similar increase in neuronal excitability and decreased ratio of E/I inputs onto these cells in 3xTg mice (**Fig. 4**). At this early time point, MECII pyramidal cells in 3xTg mice displayed no change in resting membrane potential (**Fig. 4A**, unpaired t-test, p>0.05), decreased rheobase (**Fig. 4B**, unpaired t-test, *p*=0.001) and an increased number of current-evoked action potentials (**Fig. 4C**, two-way RM ANOVA, Main Effect of Genotype: *p*=0.002) compared to WT mice. Synaptic E/I ratio was also reduced in layer II pyramidal cells of 3xTg mice (**Fig. 4D-F**, E/I frequency, Mann-Whitney, *p*=0.020; **Fig. 4G-I**, E/I amplitude, unpaired t-test, *p*=0.008), similarly skewing towards relatively more inhibitory than excitatory inputs. In contrast to the stellate cells, however, the decreased synaptic E/I ratio in MECII pyramidal cells was mainly driven by reduced excitation (**Fig. 4D**, EPSC frequency, Mann-Whitney, *p*=0.005; **Fig 4G**, EPSC amplitude, unpaired t-test, *p*=0.063). Thus, at an early pre-symptomatic time point, both MECII stellate cells and MECII pyramidal cells have increased intrinsic excitability that is concurrent with a relative enhancement of inhibition in the E/I ratio of synaptic inputs. However, each cell type appears to have distinct mechanisms driving these changes in synaptic currents, with MECII stellate cells receiving enhanced inhibitory inputs and MECII pyramidal cells receiving fewer excitatory inputs.

**Figure 4.**
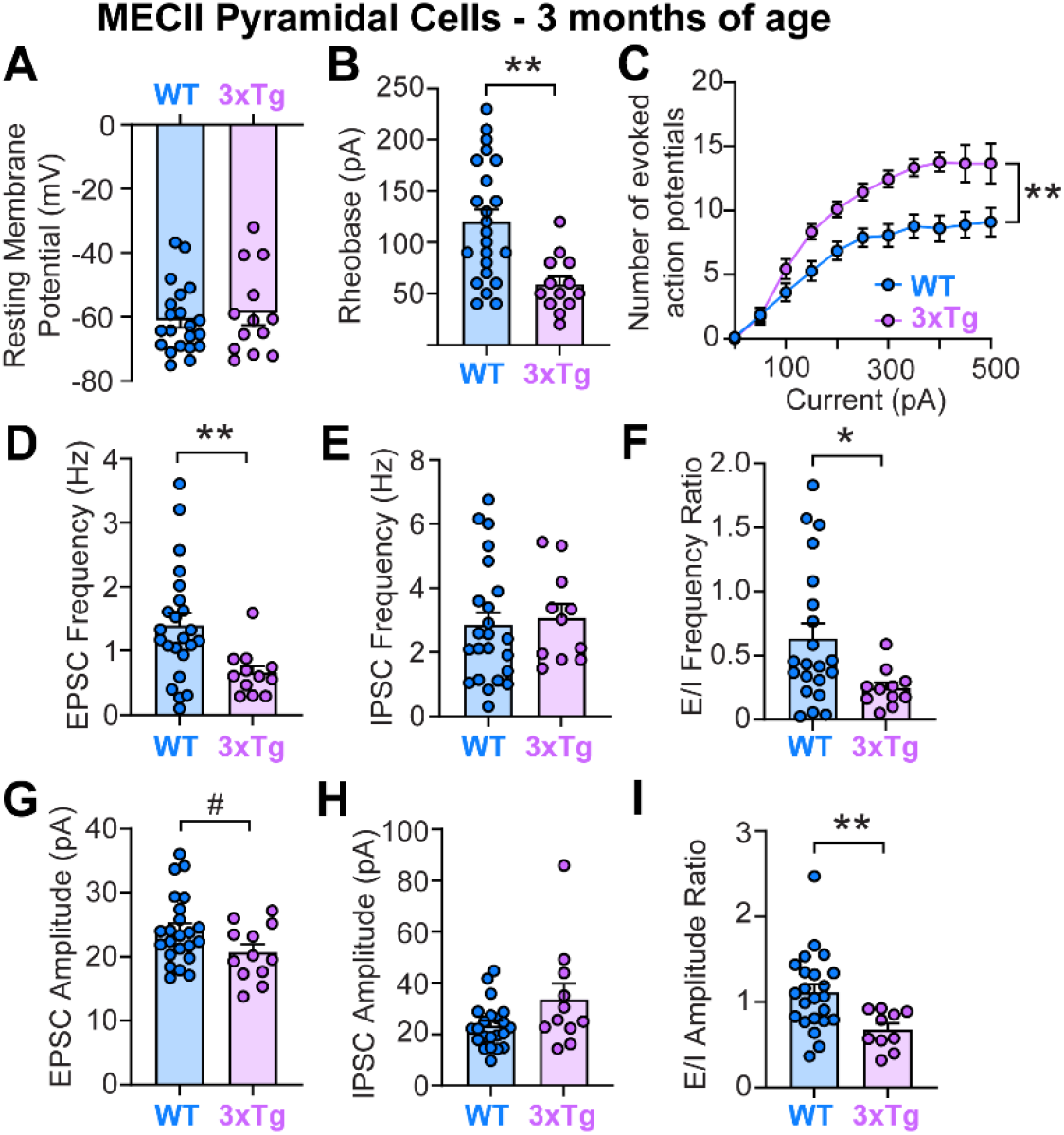
At 3 months of age, MECII pyramidal cells in 3xTg mice had increased neuronal intrinsic excitability, which was balanced by decreased synaptic excitation and decreased synaptic E/I ratio. (**A-C**) Similar to stellate cells, neuronal intrinsic excitability of MECII pyramidal cells was increased in 3xTg mice compared to controls, indicated by a reduced rheobase (**B**, unpaired t-test, *p*=0.001) and an increased number of current-evoked action potentials (**C**, n=18 cells from WT mice; n=9 cells from 3xTg mice; two-way RM ANOVA, Main Effect of Genotype: *p*=0.002). (**D-I**) Meanwhile, there was decreased synaptic excitation onto these pyramidal cells, demonstrated by a reduced EPSC frequency (**D**, Mann-Whitney test, *p*=0.005) and an overall decrease in synaptic E/I ratio both in frequency (**F**, Mann-Whitney test, *p*=0.020) and amplitude (**I**, unpaired t-test, *p*=0.008). There was also a trend for a decrease in EPSC amplitude (**G**, unpaired t-test, *p*=0.063). All other tests showed no differences between groups (*p*>0.05).

In contrast to layer II cells, MECIII neurons of 3-month-old 3xTg mice showed reduced intrinsic excitability at this early, pre-symptomatic time point (**Fig 5**). This was seen in the reduced number of current-evoked action potentials in 3xTg MECIII pyramidal cells compared to WT (**Fig. 5C**, two-way RM ANOVA, Main Effect of Genotype: *p*=0.009), despite no change in resting membrane potential or rheobase (**Fig 5A, B**). Meanwhile, 3xTg MECIII pyramidal cells were receiving more frequent synaptic inhibition (**Fig. 5E**, IPSC Frequency, unpaired t-test, *p*=0.018) and stronger excitatory inputs (**Fig. 5G**, EPSC amplitude, unpaired t-test, *p*=0.003), resulting in an overall E/I ratio that remained comparable to age-matched WT mice (**Fig. 5F**, E/I Frequency and **Fig. 5I**, E/I amplitude, *p*>0.05).

**Figure 5.**
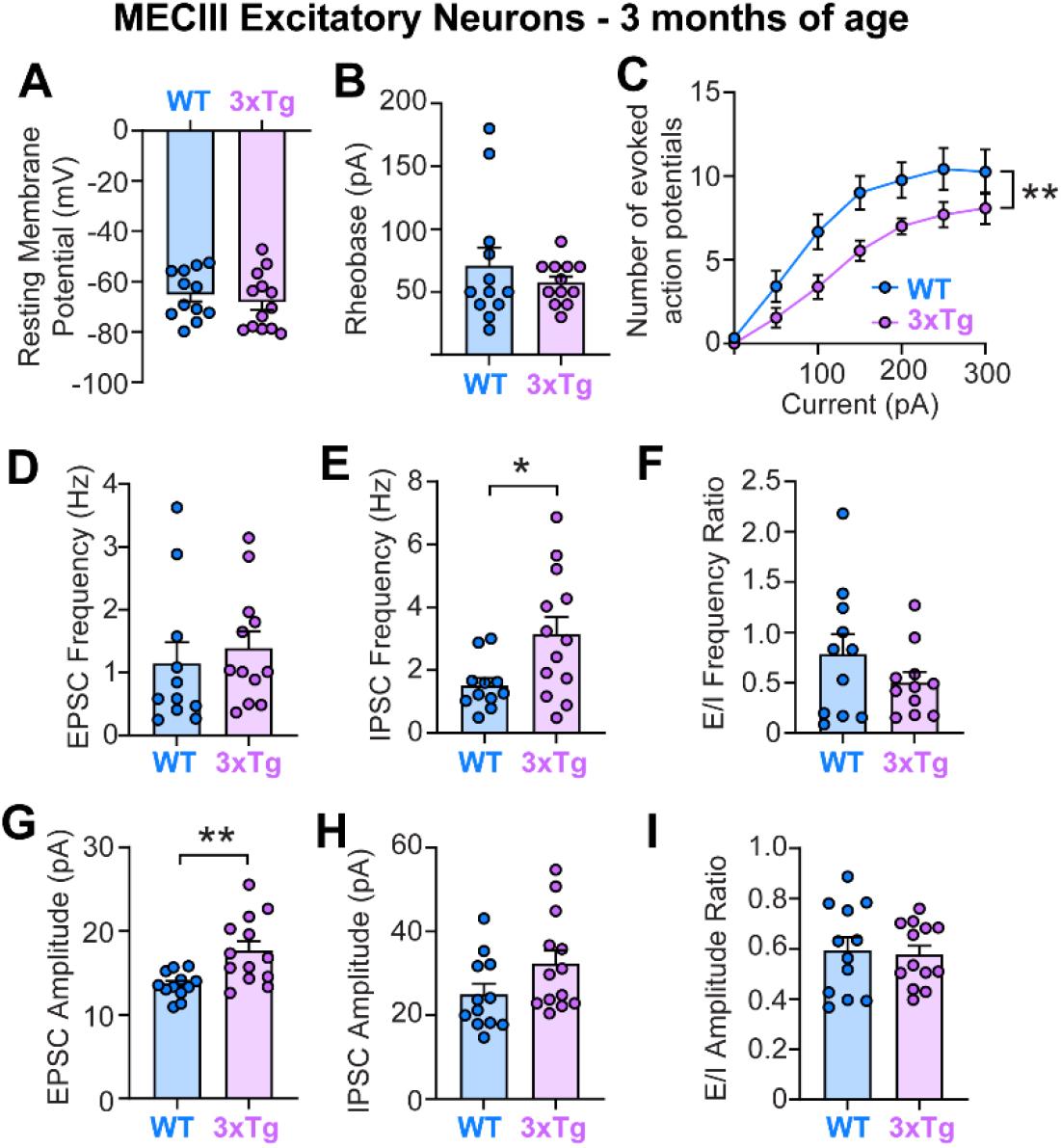
At 3 months of age, MECIII cells in 3xTg mice had decreased neuronal intrinsic excitability with no change in synaptic E/I ratio. (**A-C**) In contrast to layer II neurons, MECIII neurons showed decreased neuronal intrinsic excitability, indicated by a reduced number of current-evoked action potentials (**C**, n=12 cells from WT mice; n=13 cells from 3xTg mice; two-way RM ANOVA, Main Effect of Genotype: *p*=0.009). (**D-I**) In the same cells, there was an increase in the amplitude of EPSCs (**G**, unpaired t-test, *p*=0.003) and an increase in the frequency of IPSCs (**E**, unpaired t-test, *p*=0.018), but there were no changes in the synaptic E/I ratio in either frequency (**F**) or amplitude (**I**) of synaptic currents. All other tests showed no differences between groups (*p*>0.05).

In addition to quantifying resting membrane potential, rheobase, and the current-spike relationship, we also quantified several other intrinsic excitability measures in MECII stellate cells and MECII and MECIII pyramidal cells from 3-month-old WT and 3xTg mice (**Table 3**). In general, these additional measurements supported the main findings in **Figs. 3-5**. MECII stellate and pyramidal cells displayed early neuronal hyperexcitability (decreased spike half-width in stellate cells and increased input resistance in pyramidal cells; **Table 3**) and MECIII cells showed early hypoexcitability (a trend toward increased spike half-width, p=0.052, **Table 3**) in 3xTg mice. We also found that MECII pyramidal cells had an enlarged post-burst slow afterhyperpolarization (sAHP) which would likely reduce overall excitability (Disterhoft and Oh, 2006; Oh et al., 2010) and may serve as compensation for the early hyperexcitability seen in other measures (i.e., reduced rheobase and increased stimulated action potential firing in **Fig. 4B, C**, increased input resistance in **Table 3**).

**Table 3.** Neuronal intrinsic excitability measurements – early time point (3 m).

| **MECII Stellate cells** | **3 m WT** | **3 m 3xTg** | **p value** | **Test** |
| --- | --- | --- | --- | --- |
| Input resistance (MΩ) | 89.98 ± 8.48 | 112.6 ± 10.98 | 0.1871 | Unpaired t test |
| Spike amplitude (mV) | 85.55 ± 2.25 | 84.26 ± 1.94 | 0.7227 | Unpaired t test |
| Spike half-width (ms) | 1.88 ± 0.16 | 1.48 ± 0.07 | **\*\* 0.0101** | Mann-Whitney |
| mAHP (mV) | -6.04 ± 0.45 | -6.59 ± 0.56 | 0.5392 | Unpaired t test |
| sAHP (mV) | -0.45 ± 0.09 | -0.26 ± 0.08 | 0.1549 | Unpaired t test |
| Sag ratio | 0.41 ± 0.02 | 0.36 ± 0.02 | 0.1395 | Unpaired t test |
| **MECII Pyramidal cells** | **3 m WT** | **3 m 3xTg** | **p value** | **Test** |
| Input resistance (MΩ) | 139.9 ± 12.96 | 221.1 ± 18.50 | **\*\*\* 0.0005** | Mann-Whitney |
| Spike amplitude (mV) | 82.92 ± 2.01 | 86.24 ± 3.28 | 0.2668 | Mann-Whitney |
| Spike half-width (ms) | 1.83 ± 0.05 | 1.68 ± 0.12 | 0.1742 | Unpaired t test |
| mAHP (mV) | -1.64 ± 0.26 | -1.85 ± 0.35 | 0.6325 | Unpaired t test |
| sAHP (mV) | -0.35 ± 0.07 | -0.72 ± 0.15 | **\* 0.0181** | Unpaired t test |
| Sag ratio | 0.13 ± 0.02 | 0.11 ± 0.02 | 0.4911 | Unpaired t test |
| **MECIII cells** | **3 m WT** | **3 m 3xTg** | **p value** | **Test** |
| Input resistance (MΩ) | 257.6 ± 27.16 | 229.0 ± 20.52 | 0.4044 | Unpaired t test |
| Spike amplitude (mV) | 84.78 ± 1.90 | 86.63 ± 1.78 | 0.4852 | Unpaired t test |
| Spike half-width (ms) | 1.64 ± 0.06 | 1.92 ± 0.11 | # 0.0523 | Mann-Whitney |
| mAHP (mV) | -1.26 ± 0.27 | -1.84 ± 0.40 | 0.2455 | Unpaired t test |
| sAHP (mV) | -1.03 ± 0.24 | -0.58 ± 0.22 | 0.1801 | Unpaired t test |
| Sag ratio | 0.03 ± 0.01 | 0.06 ± 0.02 | 0.1683 | Mann-Whitney |

Together, these data demonstrate that at an early time point in the 3xTg mice, prior to the onset of MEC-hippocampus-dependent memory impairment (**Fig. 1B**), there were already intrinsic and synaptic changes in all the 3 major types of MECII and MECIII excitatory neurons. Remarkably, excitability changes at the neuronal and synaptic levels primarily moved in opposite directions, suggesting robust homeostatic regulation in this circuit that may allow it to maintain proper function. Notably, while MECII stellate and pyramidal cells underwent similar changes in the E/I ratio of synaptic currents, their mechanisms were distinct. Hyperexcitable stellate cells received relatively more inhibition while hyperexcitable pyramidal cells received less excitation, both resulting in a reduced synaptic E/I ratio. This indicates cell-type specific reorganization of synaptic inputs are occurring already at this early time point. These changes between intrinsic and synaptic excitability may result in the maintenance of normal levels of overall excitability in MECII and thus relatively normal inputs to the hippocampus early in AD, before the onset of spatial memory deficits. Regardless, these changes in neuronal and synaptic excitability were not sufficient to drive spatial memory deficits at this early time point (**Fig. 1B**).

### Development of combined intrinsic and synaptic hyperexcitability specifically in MECII stellate cells of 3xTg mice at the late, post-symptomatic time point

To test how intrinsic and synaptic excitability in MEC neurons were altered in 3xTg mice after the onset of spatial memory deficits, we again performed whole cell patch clamp recordings in MECII stellate cells of 3xTg and WT mice at 10 months of age (**Fig. 6**). In 3xTg mice, MECII stellate cells continued to have increased intrinsic excitability at this late time point, with an elevated number of current-evoked action potentials (**Fig. 6C**, two-way RM ANOVA, Main Effect of Genotype: p=0.047). However, in contrast to the early time point that showed reduced synaptic E/I ratios (**Fig. 3F, I**), at 10 months of age, MECII stellate cells in 3xTg mice had an increased synaptic E/I frequency compared to WT mice (**Fig. 6F**, E/I Frequency, unpaired t-test, *p*=0.009).

**Figure 6.**
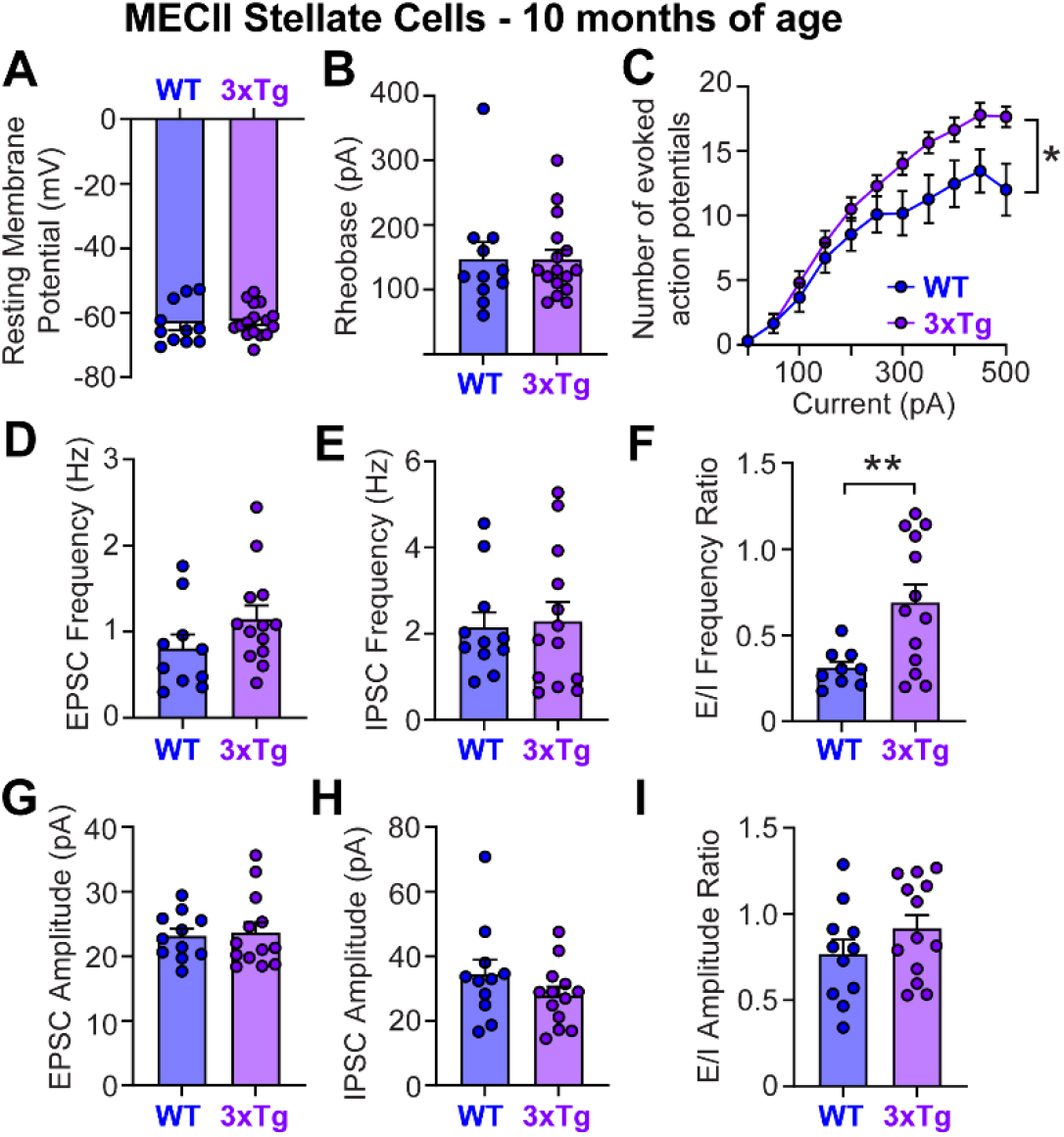
At 10 months of age, MECII stellate cells showed increases in both neuronal intrinsic excitability and synaptic E/I ratio. (**A-C**) There was an increased number of current-evoked action potentials in stellate cells of 3xTg mice (**C**, n=11 cells from WT mice; n=14 cells from 3xTg mice; two-way RM ANOVA, Main Effect of Genotype, *p*=0.048). (**D-I**) Meanwhile, the E/I frequency ratio showed a relative increase in excitatory inputs onto 3xTg stellate cells (**F**, unpaired t-test, *p*=0.009). All other tests showed no differences between groups (*p*>0.05).

Together, the heightened intrinsic excitability and synaptic E/I ratio may work synergistically to cause overall hyperexcitability in MECII stellate cells in 10-month-old 3xTg mice. Surprisingly, intrinsic and synaptic excitability were largely normalized in MECII pyramidal cells at this time point (**Fig. 7**). In MECII pyramidal neurons, we found no differences in resting membrane potential, rheobase, current-evoked action potentials, the frequency or amplitude of synaptic currents, or in the E/I balance of synaptic currents (**Fig. 7**, p>0.05 for all tests). In MECIII pyramidal cells, we found a more hyperpolarized resting membrane potential in 3xTg mice (**Fig. 8A**, unpaired t-test, *p*=0.035), but there was no change in rheobase or current-evoked firing (**Fig. 8B, C**). Meanwhile, synaptic excitation increased in these cells (**Fig. 8D**, EPSC frequency, Mann-Whitney, p=0.004), but was mostly matched by a trend toward increased inhibition (**Fig. 8E**, IPSC frequency, Mann-Whitney, p=0.073), leading to similar E/I ratios between 3xTg and WT mice (**Fig. 8F, I**). Therefore, the balance between intrinsic and synaptic excitability was generally maintained in MECIII pyramidal cells.

**Figure 7.**
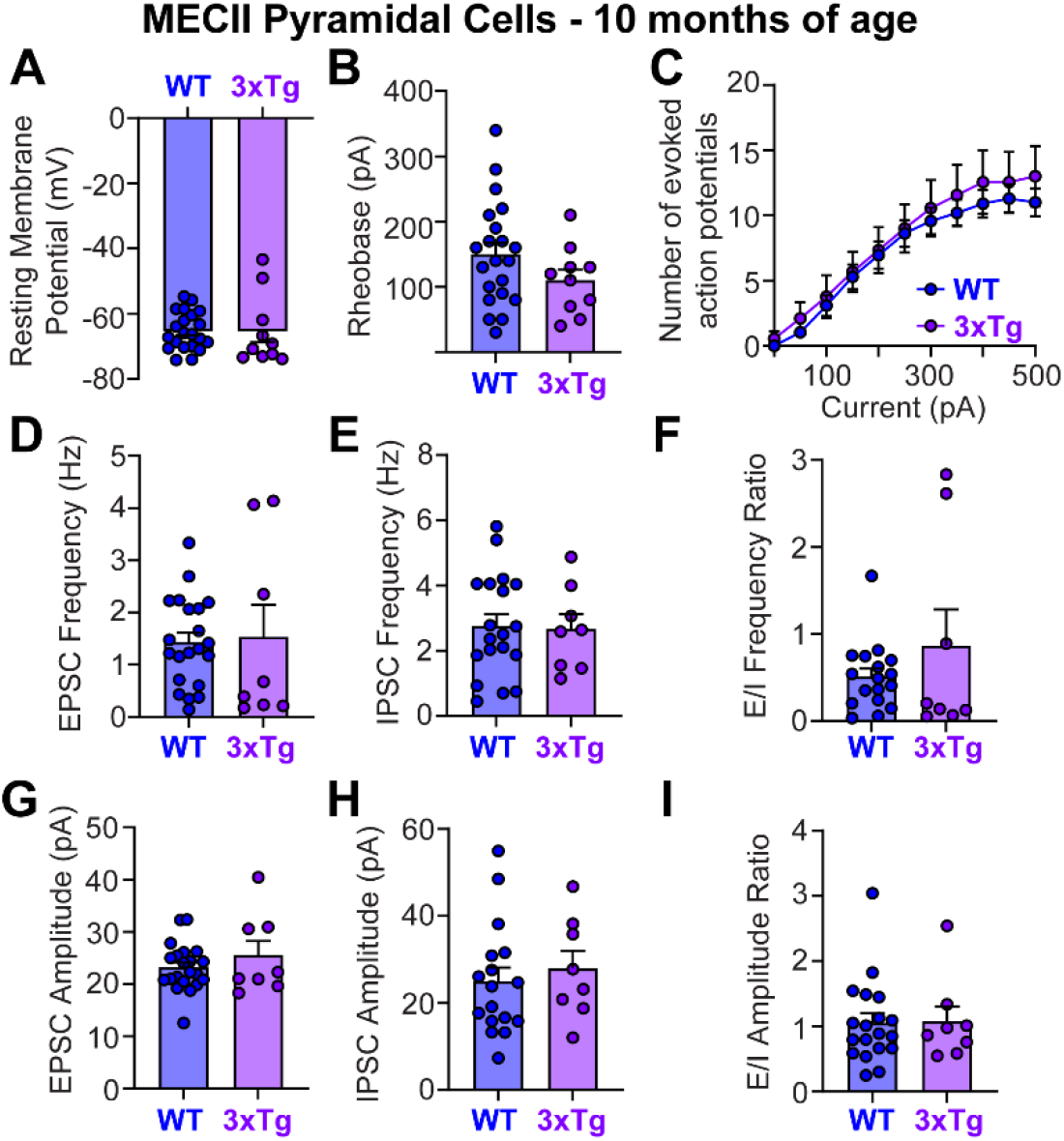
At 10 months of age, intrinsic and synaptic excitability was mostly normalized in MECII pyramidal cells in 3xTg mice. For MECII pyramidal cells, both neuronal intrinsic excitability (**A-C**) and synaptic E and I (**D-I**) in 3xTg mice were comparable to those in WT mice (all tests: *p*>0.05).

**Figure 8.**
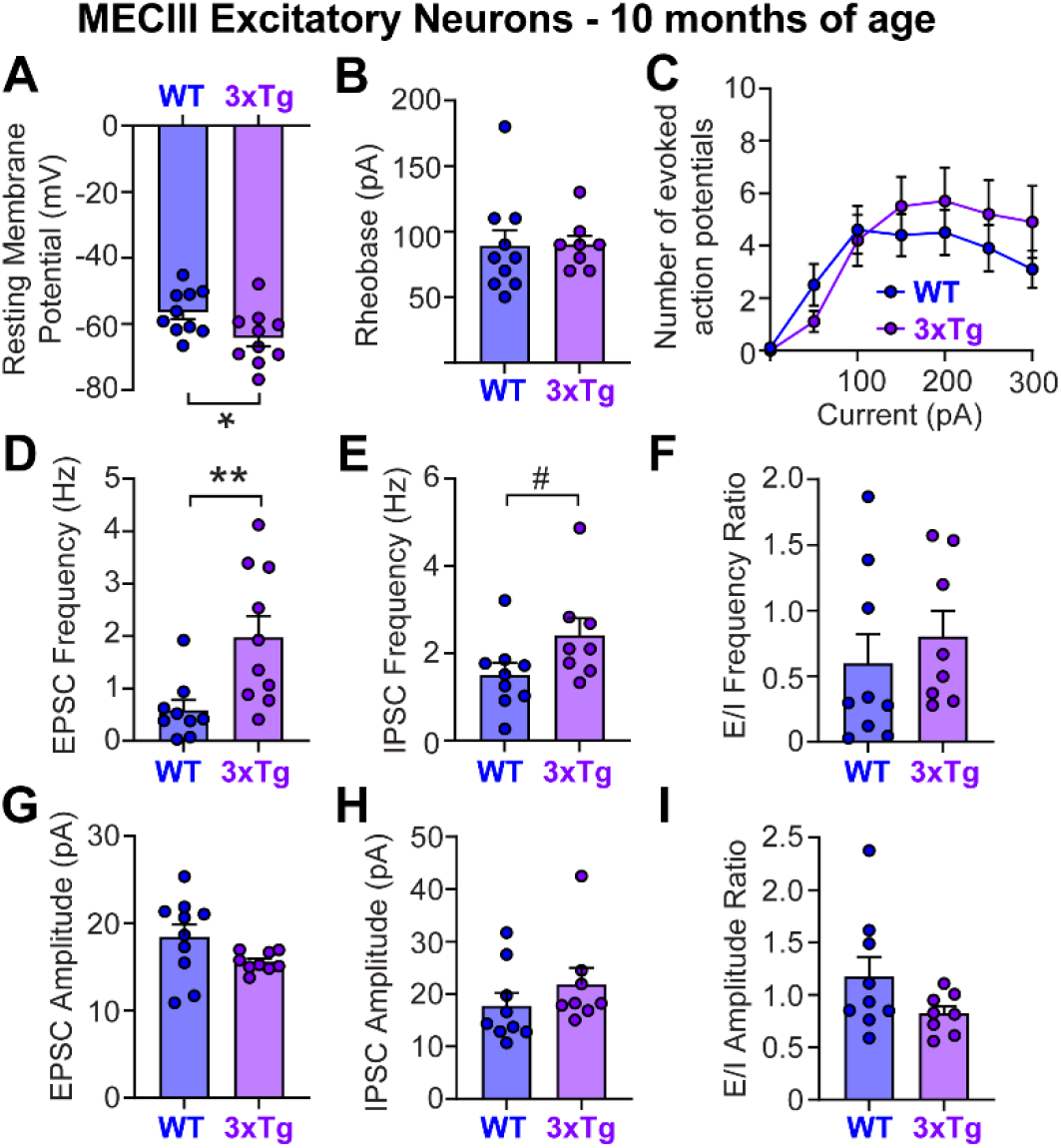
At 10 months of age, MECIII cells in 3xTg were hyperpolarized, while no change was found in synaptic E/I ratio. (**A-C**) For MECIII neurons, cells in 3xTg mice had a hyperpolarized resting membrane potential (**A**, unpaired t-test, *p*=0.035), while the rheobase and number of current-evoked of action potentials remained comparable to that in WT mice (B, C). (**D-I**) In 3xTg mice, the frequency of EPSCs was increased (**D**, Mann-Whitney test, *p*=0.004), and there was a trend for an increase in IPSC frequency (**E**, unpaired t-test, WT vs. 3xTg, *p*=0.073). However, synaptic E/I ratio did not differ between 3xTg and WT cells in either the frequency (**F**) or amplitude (**I**) of synaptic currents. All other tests showed no differences between groups (*p*>0.05).

When we looked into other intrinsic excitability properties at this late time point, significant reductions in spike half-width emerged in MECII stellate and pyramidal cells, pointing to increased neuronal excitability in 3xTg mice (**Table 4**). In MECIII pyramidal cells, while there was a hyperpolarized resting membrane potential (**Fig. 8A**), there was also a significantly reduced mAHP and a trend toward a reduced sAHP (**Table 4**) pointing to increased excitability (Disterhoft and Oh, 2006; Oh et al., 2010). This may reflect homeostatic compensation between the hyperpolarized membrane potential and the AHP amplitudes that may allow MECIII pyramidal cells to maintain relatively normal firing rates in 3xTg mice at 10 months of age.

**Table 4.** Neuronal intrinsic excitability measurements – late time point (10 m).

| **MECII Stellate cells** | **10 m WT** | **10 m 3xTg** | **p value** | **Test** |
| --- | --- | --- | --- | --- |
| Input resistance (MΩ) | 78.59 ± 8.57 | 79.13 ± 5.95 | 0.9581 | Unpaired t test |
| Spike amplitude (mV) | 85.65 ± 1.09 | 81.28 ± 1.75 | **\* 0.0444** | Unpaired t test |
| Spike half-width (ms) | 1.38 ± 0.03 | 1.28 ± 0.04 | **\* 0.0069** | Mann-Whitney |
| mAHP (mV) | -7.99 ± 0.53 | -8.14 ± 0.46 | 0.5038 | Mann-Whitney |
| sAHP (mV) | -0.34 ± 0.05 | -0.30 ± 0.04 | 0.5404 | Unpaired t test |
| Sag ratio | 0.42 ± 0.02 | 0.42 ± 0.01 | 0.4510 | Mann-Whitney |
| **MECII Pyramidal cells** | **10 m WT** | **10 m 3xTg** | **p value** | **Test** |
| Input resistance (MΩ) | 158.4 ± 21.74 | 155.9 ± 16.56 | 0.3927 | Mann-Whitney |
| Spike amplitude (mV) | 75.82 ± 2.87 | 81.29 ± 2.37 | 0.2866 | Mann-Whitney |
| Spike half-width (ms) | 1.84 ± 0.10 | 1.38 ± 0.05 | **\*\*\* 0.0005** | Mann-Whitney |
| mAHP (mV) | -1.49 ± 0.41 | -1.38 ± 0.48 | 0.6224 | Mann-Whitney |
| sAHP (mV) | -0.45 ± 0.12 | -0.28 ± 0.09 | 0.9332 | Mann-Whitney |
| Sag ratio | 0.11 ± 0.02 | 0.16 ± 0.04 | 0.1244 | Mann-Whitney |
| **MECIII cells** | **10 m WT** | **10 m 3xTg** | **p value** | **Test** |
| Input resistance (MΩ) | 267.3 ± 15.71 | 225.5 ± 30.65 | 0.2408 | Unpaired t test |
| Spike amplitude (mV) | 64.08 ± 1.86 | 67.35 ± 2.72 | 0.3350 | Unpaired t test |
| Spike half-width (ms) | 2.09 ± 0.17 | 1.92 ± 0.11 | 0.4054 | Unpaired t test |
| mAHP (mV) | -2.41 ± 0.53 | -0.56 ± 0.25 | **\*\* 0.0048** | Unpaired t test |
| sAHP (mV) | -0.38 ± 0.09 | -0.12 ± 0.09 | # 0.0588 | Unpaired t test |
| Sag ratio | 0.15 ± 0.03 | 0.08 ± 0.04 | 0.1903 | Mann-Whitney |

Taken together, these results highlight a cell-type specific shift towards overall hyperexcitability in MECII stellate cells at the late, post-symptomatic time point in the 3xTg mouse model of AD pathology. Notably, the increased synaptic inhibition that was observed at 3 months of age was not present at 10 months of age, and instead the E/I ratio switched from a shift toward inhibition to a shift toward excitation during the emergence of spatial memory impairments in 3xTg mice. These intrinsic and synaptic alterations drive MECII stellate cell hyperexcitability at this late time point, which may drive changes in the downstream hippocampal activity and the emergence of memory impairments. Importantly, this overall hyperexcitability, with both intrinsic and synaptic changes strongly driving increased excitability, was only observed in MECII stellate cells at the late, post-symptomatic time point. In contrast, at the post-symptomatic time point, we found minimal changes in excitability of MECII pyramidal and MECIII excitatory neurons, suggesting that these neurons are unlikely to drive memory impairments in 3xTg mice.

## DISCUSSION

Here, we examined neuronal intrinsic excitability and synaptic activity in three major subtypes of MEC excitatory cells during the progression of memory impairments in the 3xTg mouse model of AD pathology (**Fig. 9**). We found that prior to the onset of memory impairments, early neuronal and synaptic changes were present in all three cell types, but intrinsic changes were generally balanced by opposing shifts in synaptic inputs. This suggests that homeostatic mechanisms were in place, presumably maintaining overall levels of excitability in MEC at this early, pre-symptomatic time point. Conversely, after the onset of memory impairments, we found that MECII stellate cells were especially hyperexcitable, with both increased intrinsic hyperexcitability and increased synaptic excitation relative to inhibition. This data suggests that there are compensatory mechanisms in place that enhance the inhibitory inputs onto these cells at the early time point, but that these mechanisms become compromised during the progression of memory impairments, exacerbating the increased excitability in MECII stellate cells. This increased excitability in stellate cells may contribute to memory impairments as stellate cells are necessary for spatial memory (Tennant et al., 2018) and increased excitability in these cells is sufficient to drive hippocampal remapping and memory impairments in control mice (Kanter et al., 2017). Thus, the increased excitability we observed in MECII stellate cells at the 10 month time point is a potential mechanism that may directly contribute to spatial memory impairment in this 3xTg model of AD pathology.

**Figure 9.**
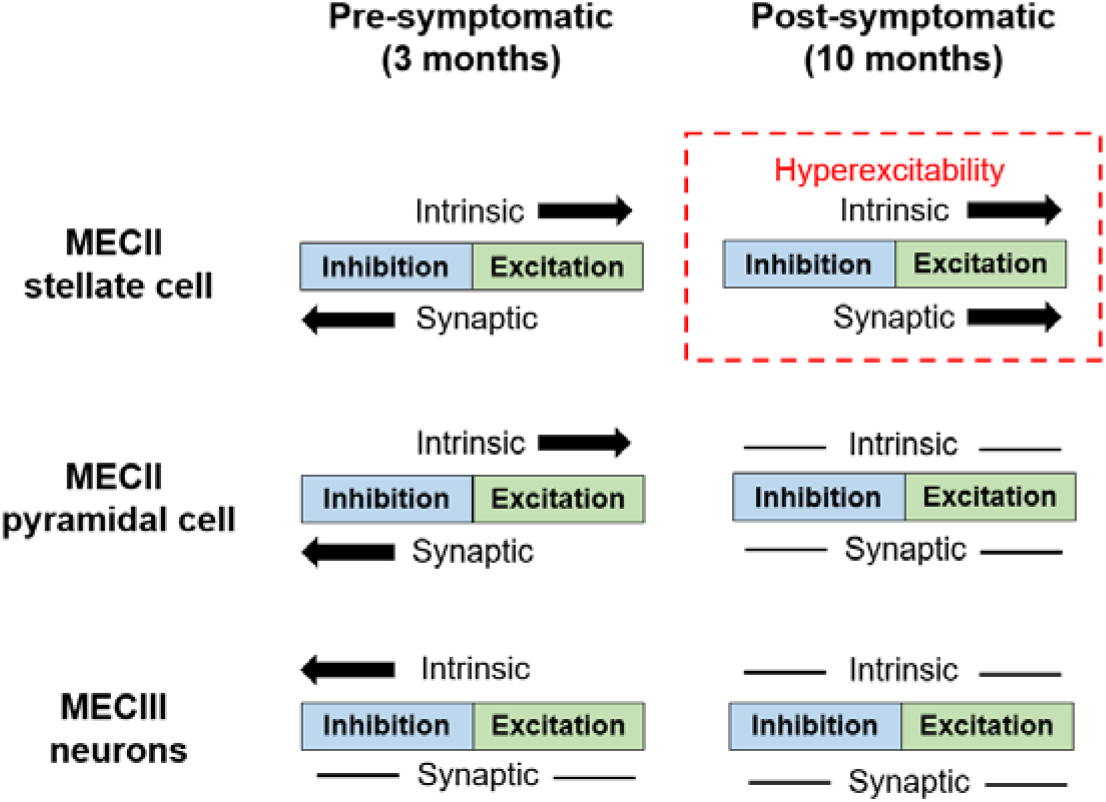
Summary of intrinsic and synaptic changes in excitability across MECII stellate cells, MECII pyramidal cells, and MECIII neurons in 3xTg mice. Progressive changes in intrinsic and synaptic excitability in MECII stellate cells may lead to cell-type specific hyperexcitability in 3xTg mice. At the early, pre-symptomatic time point (3 months), neuronal intrinsic excitability changes in young 3xTg mice are matched by alterations in synaptic excitation and inhibition in a way that helps to maintain overall excitability in MEC neurons. At the late, post-symptomatic time point (10 months), while the intrinsic and synaptic excitability in MECII pyramidal cells and MECIII neurons have largely normalized, MECII stellate cells in 3xTg mice remain hyperexcitable both at the intrinsic and synaptic levels. This loss of homeostatic balance may cause local network hyperexcitability, disrupt downstream hippocampus activity, and lead to deficits in spatial memory.

Neuronal and synaptic changes have been primarily studied in the hippocampus in mouse models of AD pathology. Previous studies have found reduced efficacy of synaptic transmission in the hippocampus (Oddo et al., 2003; Palop and Mucke, 2010; Booth et al., 2016b; Forner et al., 2017; Chen et al., 2018), and several studies have found increased intrinsic excitability across mouse models of AD pathology in CA1 (Brown et al., 2011; Davis et al., 2014; Kerrigan et al., 2014; Frazzini et al., 2016) and the dentate gyrus (Minkeviciene et al., 2009; Jiang et al., 2021). However, AD-related neurophysiological changes in the MEC are less well-defined. In symptomatic Tg2576 mice, a model of mutant APP overexpression, MECII stellate cells displayed a slight increase in firing frequency during mild depolarization but otherwise normal excitability during higher stimulations (Marcantoni et al., 2014). In these mice, perforant path input was also compromised both structurally and functionally before detectable plaque deposition (Jacobsen et al., 2006), suggesting early changes in entorhinal inputs to hippocampus.

Furthermore, in the McGill-R-Thy1-APP transgenic rat model with progressive plaque pathology, stellate cells were more excitable before plaque deposition, while the electrophysiology properties of other types of MECII principal cells remained unaltered (Heggland et al., 2019). In accordance with these cellular changes, impairments in grid cell coding were found in mouse models of AD pathology (Fu et al., 2017; Ying et al., 2022; Igarashi, 2023), even before the onset of spatial memory deficits (Jun et al., 2020). In adult humans at genetic risk for AD, there were reduced grid-cell-like representations as well (Kunz et al., 2015). Together, these studies point to early changes of neuronal excitability in stellate cells that may further affect perforant path input to the hippocampus.

By recording intrinsic and synaptic properties in the same neurons, we have indeed observed early intrinsic and synaptic changes in the MEC of 3xTg mice, before the onset of memory impairments, which typically occur around 4-6 months of age in this model (Billings et al., 2005; Davis et al., 2013; Belfiore et al., 2019; Creighton et al., 2019). At the pre-symptomatic time point, intrinsic and synaptic excitability changes in MECII occurred in opposing directions, which likely helps maintain normal levels of MECII output.

Interestingly, different subtypes of MECII excitatory neurons recruited distinct synaptic mechanisms in balancing intrinsic excitability changes at 3 months of age. To balance increased intrinsic excitability in 3xTg mice, stellate cells received increased inhibition while pyramidal cells received reduced excitation compared to WT mice. This distinction is particularly interesting considering that this enhanced inhibition onto stellate cells was lost at the late time point (10 months) when those cells were still hyperexcitable, exacerbating overall network hyperactivity. Increased inhibitory synaptic input onto stellate cells at the pre-symptomatic time point may originate from pathology-related neurophysiological alterations of interneurons themselves or through a homeostatic circuit mechanism. At the late time point, the balancing effect of increased synaptic inhibition was lost, possibly due to impaired inhibitory synaptic transmission or interneuron dysfunction, which has been observed in the hippocampus (Palop et al., 2007; Leung et al., 2012; Hazra et al., 2013; Schmid et al., 2016; Prince et al., 2021), parietal cortex (Verret et al., 2012; Chen et al., 2018), and lateral entorhinal cortex (Petrache et al., 2019) in several mouse models of AD pathology. In MEC, inhibition is mainly provided by parvalbumin (PV)- and cholecystokinin (CCK)-expressing interneurons at the perisomatic regions and somatostatin (SOM)-expressing interneurons at dendrites (Varga et al., 2010; Fuchs et al., 2016; Martínez et al., 2017; Fernandez et al., 2022). While PV interneurons innervate layer II stellate and pyramidal cells similarly (Varga et al., 2010; Armstrong et al., 2016; Fernandez et al., 2022), CCK cells preferentially target pyramidal cells (Varga et al., 2010; Fuchs et al., 2016). PV interneurons provide strong somatic inhibition and play an important role in tuning the excitability of principal cells (Freund and Katona, 2007). Moreover, PV cell dysfunction has been characterized in AD mouse models and linked to learning and memory deficits (Verret et al., 2012; Chen et al., 2018; Petrache et al., 2019). Therefore, it is likely that time-dependent changes in the number, function, and synaptic innervation of PV interneurons may drive the switch in synaptic inhibition from high to low in stellate cells. Further research on the intrinsic excitability of PV and other interneurons are necessary to elucidate the role of inhibition on tuning network excitability in the MEC across the progression of AD.

AD is characterized by two major pathological hallmarks: the accumulation of extracellular amyloid beta (Aβ) plaques and intracellular neurofibrillary tangles made of hyperphosphorylated tau. Both of these pathologies seem to affect neuronal excitability and circuit dynamics (Harris et al., 2020). In general, studies have found that soluble Aβ oligomers in early AD are causally linked to neuronal and circuit hyperexcitability (Brorson et al., 1995; Ye et al., 2003; Busche et al., 2008, 2012; Zott et al., 2019). On the contrary, tau pathology has generally exhibited an opposing effect (Menkes-Caspi et al., 2015; Fu et al., 2017; Busche et al., 2019; Marinković et al., 2019). Importantly, amyloid and tau pathologies have complex interactions with each other that mediate neural activity (Roberson et al., 2007, 2011; Ittner et al., 2010; Busche et al., 2019). Therefore, to capture the combined effects of pathological Aβ and tau on the effect of neuronal excitability and synaptic transmission, we utilized the 3xTg mouse model of AD pathology which has both amyloid and tau mutations (Oddo et al., 2003; Billings et al., 2005). In this model, Aβ production is enhanced by the APP Swedish mutation (Haass et al., 1995), which increases cleavage of APP by β-secretase, and the PS1_M146V_ mutation, which increases production of Aβ_42_ (Jankowsky et al., 2004), the Aβ variant highly linked with amyloid aggregation. In addition, tau pathology is driven by the MAPT_P301L_ transgene that induces tau hyperphosphorylation and neurofibrillary tangles. In 3xTg mice, intracellular Aβ immunoreactivity is apparent between 3 and 4 months of age in the neocortex and is correlated with the emergence of cognitive deficits, while extracellular plaques appear around 6 months of age in the frontal cortex and become evident in other cortical regions and hippocampus by 12 months (Oddo et al., 2003; Billings et al., 2005). In the entorhinal cortex, extracellular plaques also emerge around 12 months of age (Belfiore et al., 2019). In addition, phosphorylated tau has been detected in the hippocampus and cortex as early as 4 months of age in 3xTg mice, and is highly prominent by 12 months of age (Belfiore et al., 2019; Javonillo et al., 2022).

Since soluble Aβ is toxic (Oddo et al., 2003; Busche et al., 2012; Mucke and Selkoe, 2012; Xu et al., 2015; Busche and Konnerth, 2016), initial accumulation of Aβ oligomers in 3-month-old 3xTg mice may drive the observed changes in neuronal intrinsic excitability and synaptic transmission at this early stage. However neuronal and synaptic changes maintained a homeostatic balance at this time point, and MEC-dependent memory remained intact. Interestingly, MECII stellate and pyramidal cells displayed hyperexcitability while MECIII cells showed hypoexcitability, indicating potential differential accumulation and/or effects of intracellular Aβ to these cell types. Surprisingly, at the late time point (10 months), neuronal excitability became largely normalized in MECII pyramidal neurons and in MECIII excitatory neurons. This could be the result of opposing effects of Aβ and tau pathologies, with distinct temporal onsets, as hyperactivity appears first in preclinical stages of AD and hypoactivity in later phases of the disease (Dickerson et al., 2005; Busche and Konnerth, 2016; Harris et al., 2020). Here, we may have captured this switch during the evolving pathological changes in the MEC, leading to a temporarily normalized excitability profile in MECII pyramidal cells and MECIII excitatory neurons. Thus, at an even more advanced age, MEC neurons may further shift towards hypoexcitability, similar to what is observed in human patients. Regardless, the predominant circuit change in MEC that corresponded with the onset of memory impairments was the overall increased excitability in both the intrinsic and synaptic properties of MECII stellate cells.

Overall, the time-dependent and cell-type specific changes in the three major types of excitatory neurons in the MEC indicate complex changes associated with AD pathology. Understanding this temporal progression in distinct cell types will enable us to develop early treatments and interventions to slow down or halt the development of cognitive dysfunction in this disease. Further studies using cell-type specific optogenetic, chemogenetic, or pharmacogenetic manipulations will help elucidate any causal relationship between cell-type specific abnormal neural activity in the MEC and the progression of memory deficits in Alzheimer’s disease.

## ACKNOWLEDGEMENTS

We would like to thank all members of the Shuman and Cai labs for their comments and feedback throughout this project. This work was supported by NIH grants RF1AG072497 (TS), R01NS116357 (TS), F32NS116416 (ZCW), F31AG069496 (LV), R01MH120162 (DJC), DP2MH122399 (DJC), and an American Epilepsy Society Predoctoral Fellowship (YF).

